# Inferring processes of community assembly from macroscopic patterns: the case for inclusive and mechanistic approaches

**DOI:** 10.1101/195008

**Authors:** Mikael Pontarp, Åke Brännström, Owen L Petchey

## Abstract

Statistical techniques exist for inferring community assembly processes from community patterns. Habitat filtering, competition, and biogeographical effects have, for example, been inferred from signals in phenotypic and phylogenetic data. The usefulness of current inference techniques is, however, debated as the causal link between process and pattern is often lacking and processes known to be important are ignored. Here, we revisit current knowledge on community assembly across scales and, in line with several reviews that have outlined the features and challenges associated with current inference techniques, we identify a discrepancy between features of real communities and current inference techniques. We argue, that mechanistic eco-evolutionary models in combination with novel model fitting and model evaluation techniques can provide avenues for more accurate, reliable and inclusive inference. To exemplify, we implement a trait-based and spatially explicit dynamic eco-evolutionary model and discuss steps of model modification, fitting, and evaluation as an iterative approach enabling inference from diverse data sources. This suggested approach can be computationally intensive, and model fitting and parameter estimation can be challenging. We discuss optimization of model implementation, data requirements and availability, and Approximate Bayesian Computation (ABC) as potential solutions to challenges that may arise in our quest for better inference techniques.

## Introduction

Community assembly processes are difficult to observe directly in the field and revealing processes via manipulative experiments is not always feasible. Consequently, there is a considerable need to infer processes from observations, such as trait-distributions, species distributions, abundances, and phylogenies (Emerson and Gillespie 2008; Cavender-Bares et al. 2009; Vamosi et al. 2009; Cadotte et al. 2010; Pausas and Verdu 2010; Mouquet et al. 2012). As an example, the co-occurrence of species having similar niches (high phenotypic clustering) or dissimilar niches (low phenotypic clustering, also termed overdispersion) may reflect habitat filtering or ecological interactions, respectively (Webb et al. 2002). Other common methods quantify the correlation between species occurrence and abiotic factors (Legendre et al. 1997) or distance between habitats (Borcard et al. 1992) to infer habitat filtering or geographical contingencies.

Despite their widespread and frequent use, current assembly process inference techniques have limitations. Most methods rely on statistical models for one or a few processes, although it is well known that community assembly occurs via multiple processes (Ackerly et al. 2006; Ricklefs 2007; Leibold et al. 2010). Patterns observed in nature may be consistent with multiple explanations (Vellend 2010) and current techniques may thus fail to provide accurate inference, particularly if evolutionary processes and trophic interactions are poorly integrated (Emerson and Gillespie 2008; Pausas and Verdu 2010; Pontarp and Petchey 2016). Fundamental assumptions (e.g. that competition will result in overdispersed trait distributions), on which current inference techniques often rely, have also been questioned (Mayfield and Levine 2010). Such challenges and shortcomings (as well as advantages) of existing inference techniques are covered in several reviews (Emerson and Gillespie 2008; Cavender-Bares et al. 2009; Vamosi et al. 2009; Cadotte et al. 2010; Pausas and Verdu 2010; Mouquet et al. 2012; Adler et al. 2013). Our aim is thus not to review current inference techniques, though we outline the most relevant features of some of the common ones (Tables 1-2 and Online Appendix 1). Instead, we argue that a synthesis of existing modeling frameworks and statistical techniques have the potential to transform the practice of inference of process from pattern in ecology and evolutionary biology.

**Table 1.**
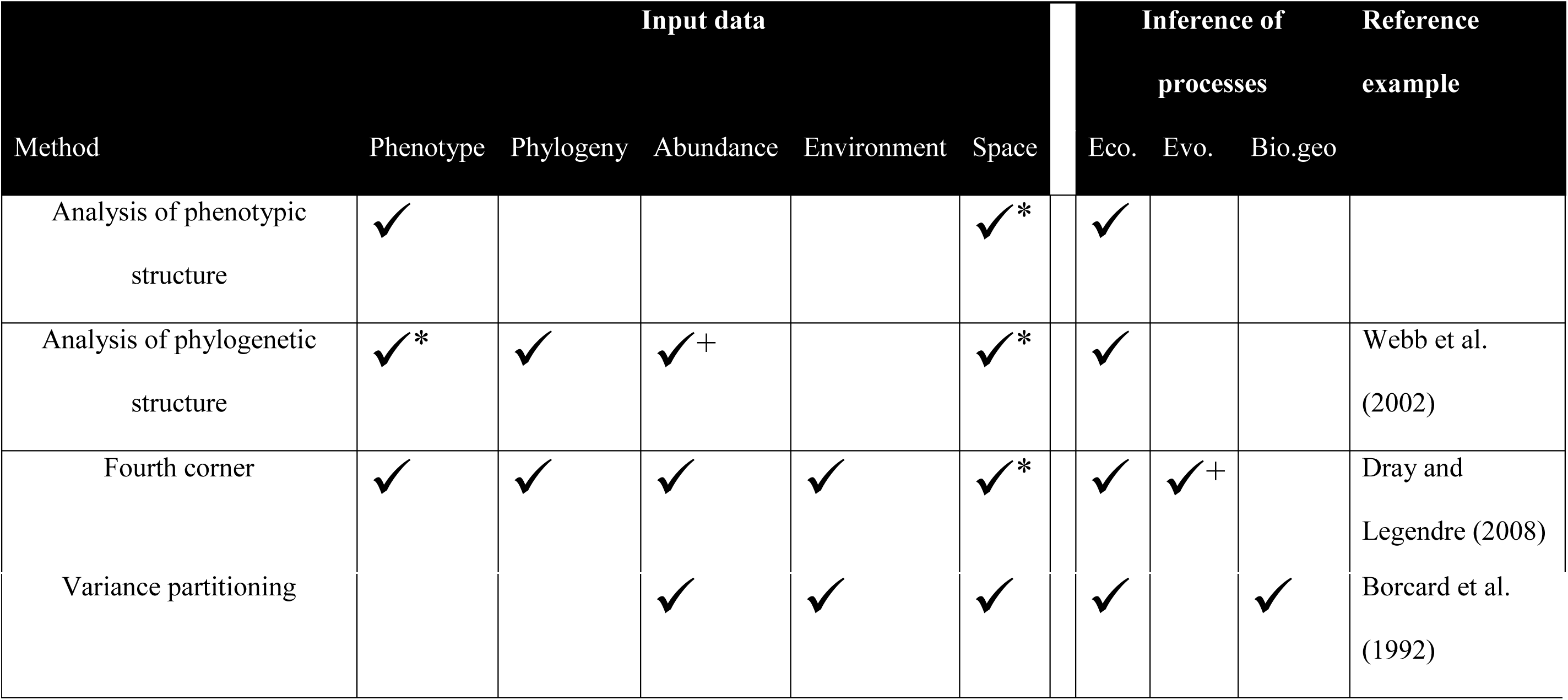
Methods that infer assembly processes from community patterns (column 1) and the information that they consider (column 2-5). Tick mark denotes which data/processes are considered explicitly. Superscript denotes data/processes that are implicitly considered (*) or included in extensions of the basic method (+). “Eco” processes include habitat filtering and competitive exclusion; “Evo” include the evolution of phenotypes, for example via character displacement; “Bio. geog.” includes dispersal limitation.

**Table 2.** Challenges and limitations associated with current inference techniques and methods categorized into three overarching categories (column 1).

| Challenge category | Number | Challenge/ Limitation | Description | Source example |
| --- | --- | --- | --- | --- |
| Basic assumptions of the methods | i | Identifying traits can be used as a proxy for niche. | Phenotypic inference techniques assume that trait(s) can be used as a proxy for niche. Identifying the traits that define an organism's niche and thus drive the assembly processes can, however, be difficult and must be supported by expert knowledge of organisms' natural histories. Appropriate weighting of traits can also be difficult to establish. | Petchey and Gaston (2006) |
|  | ii | Identifying the relationship between niche, traits, and phylogeny. | Phylogenetic inference techniques assume a mapping between relatedness and niche. The interpretation of phylogenetic patterns is contingent on trait evolution and the distribution of traits over the phylogeny. The degree of community clustering given by a certain process will, in other words, be contingent on whether the niche is conserved or labile. | Wiens et al. (2010) |
|  | iii | Assuming a fixed species pool and no evolution or no explicit space | The current theory is mainly focused on local-community assembly from a fixed regional species pool and assumes that dispersal is not limiting. This implies that local community composition results only from local processes (e.g., habitat filtering and competition). | Pausas and Verdu (2010) |
|  | iv | Assuming that only similar species exclude each other due to competition. | A current paradigm in ecology states that species that are closely related, share traits and thus also have similar niches cannot co-exist due to competitive exclusion. Recent studies have, however, showed that competition can sometimes eliminate more different and less related species. | Mayfield and Levine (2010) |

Table 2 continued
| Challenge category | Number | Challenge/ Limitation | Description | Source example |
| --- | --- | --- | --- | --- |
| Method specificity and scale-dependent processes | v | Single-process inference | Most methods only allow for inference on the net effect of processes and do not partition the relative importance of ecological, evolutionary, and spatial processes acting in concert. | Petchey (2007) |
|  | vi | Evolution is ignored | Evolutionary contingencies are largely ignored in current inference techniques. This is limiting, in particular when analyzing phylogenetic patterns. | Emerson and Gillespie (2008) |
|  | vii | Scale-dependent processes | Different traits may be affected by different processes at different temporal and spatial scales. | Trisos et al. (2014) |
| Lack of mechanistic links between process and | viii | Unknown mechanisms | Methods that take environmental factors and explicit spatial components into account (e.g., variance partitioning) are mainly phenomenological or statistical. A mechanistic understanding of the causal link between pattern and process is thus often lacking. | Gotelli et al. (2009) |
|  | ix | Trophic interactions are often ignored | Even though it is well known that trophic interactions can structure communities, trophic processes are largely ignored when patterns are interpreted. | Mouquet et al. (2012) |

We set the stage by reviewing current knowledge on the diversity of community assembly processes that have been termed the “black box of community ecology” by Vellend (2010). With community assembly processes across spatiotemporal scales in mind, we thereafter emphasize the need for more holistic assembly-process inference. Such transformation, already underway, involves more mechanistic and complex models of community assembly. We highlight specific components of such a transformation including mechanistic modeling, parameter estimation, and model selection (see also Csillery et al. 2010; van der Plas et al. 2015; Cabral et al. 2017).

In a concrete example, we present a trait-based and spatially explicit dynamic eco-evolutionary community model that includes various processes ranging from intra- and interspecific competition, trophic interactions, dispersal as well as trait evolution within trophic levels and co-evolution among trophic levels. We choose to implement our model as a differential equations (Kot 2001) and matrix model which allows for tractable computational cost with the flexibility to initiate simulations with different conditions including or excluding particular processes. We then use this model to illustrate how steps of model modification (including or excluding processes) can be customized for a range of data sources including time series of population abundance, phylogenies, trait distributions, and spatial species distribution data. Furthermore, we argue that an iterative approach of model modification, model fitting, and model evaluation can answer calls for novel inference techniques.

Using complex models and sophisticated parameter estimation techniques come with challenges associated with data requirements, computational costs, and statistical issues such as model fitting and selection. Many of these challenges are already recognized in other fields such as ecological forecasting and data assimilation (Luo et al. 2011; Niu et al. 2014). Here we synthesize and assess them for the purposes of process inference. We envision that attempts to overcome such challenges, through a combination of data collection, experimental work and a well-defined inference framework will be a worthwhile endeavor on the road towards mechanistic inference of multiple community assembly processes acting in concert.

## Processes of eco-evolutionary metacommunity assembly

Before delving into the technical details, it is important to recognize the complex nature of community assembly across spatiotemporal scales. On *short temporal and small geographical scales in communities with no trophic interactions*, habitat filtering (Wilson et al. 1999) and limiting similarity (MacArthur and Levins 1967) have been viewed as the dominating assembly processes (Fig. 1). The abiotic environment may filter the community such that only species with traits that facilitate their survival within particular environments (e.g. temperature or levels of precipitation) can persist. Habitat filtering will thus cause a local community to become phenotypically clustered and, if traits are phylogenetically conserved (Blomberg et al. 2003), also phylogenetically clustered. On the contrary, but not mutually exclusive, competition can drive community overdispersion as superior competitors outcompete inferior ones. With this being said, the paradigm of habitat filtering and competition driving community clustering and overdispersion have been challenged by recent studies that show trait convergence among competing species (Mayfield and Levine 2010; Godoy et al. 2014; Kraft et al. 2015).

**Fig. 1.**
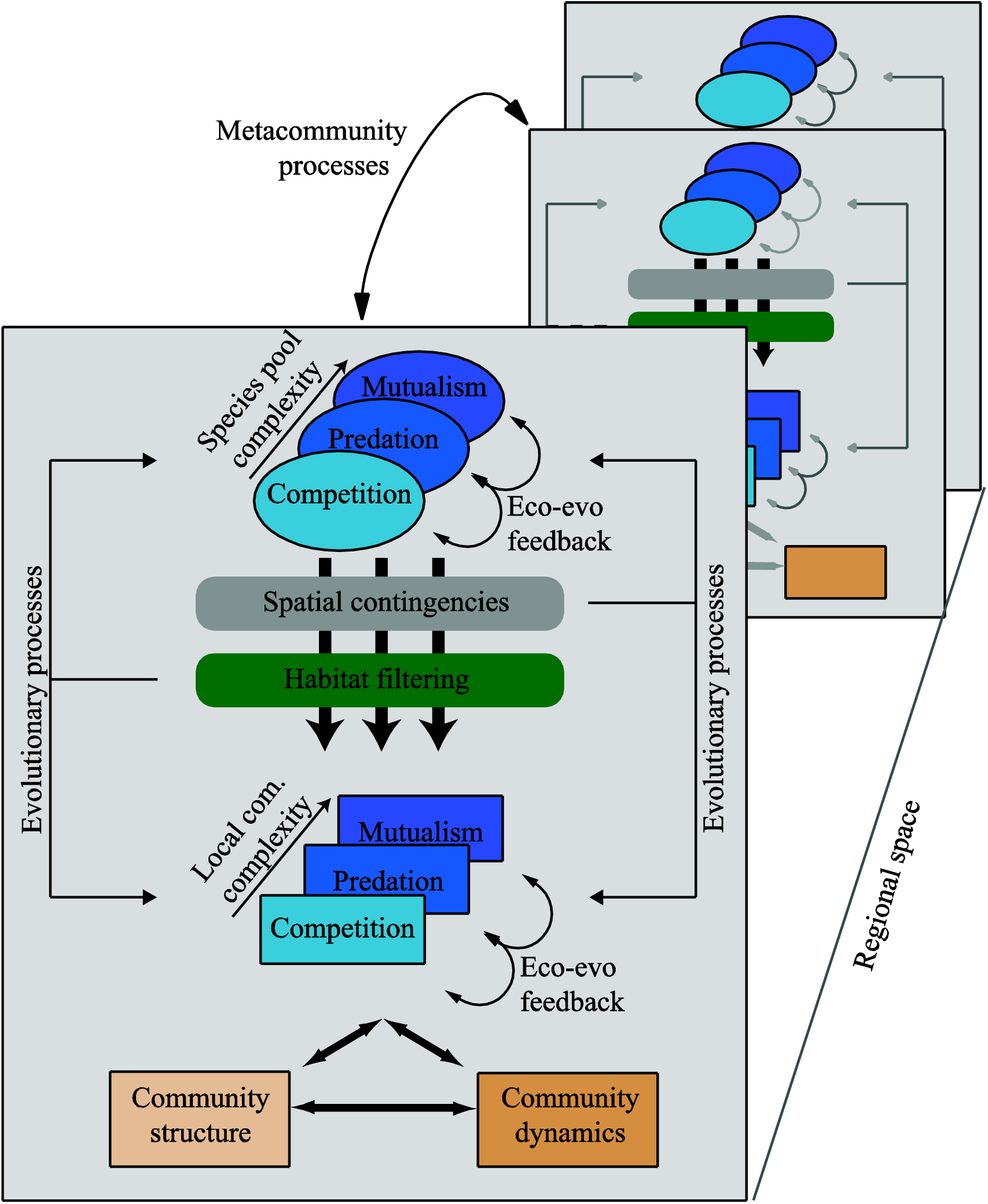
Illustration of the link between community assembly processes on different spatiotemporal scales and community structure/dynamics. A common view of local community assembly is based on a competitive community (light blue rectangle) which is assembled from a local species pool (light blue ellipse) through habitat filtering and ecological interactions such as competition. The assembly processes dictate community structure and dynamics. It is, however, also well known that local spatial contingencies (gray rectangle) and metacommunity processes dictate community assembly. Furthermore, natural communities often contain predation and mutualistic interactions adding to the community complexity of any species pool and local community. Moreover, spatial contingencies and habitat filtering often feedback into local community and species pool structure through evolutionary processes. Similarly, eco-evolutionary feedback processes affect both local species and the species pool.

In *trophic communities*, both empirical (Alto et al. 2012) and theoretical (Pontarp and Petchey 2016) studies show that trait-dependent trophic interactions also structure communities (Fig. 1). When correlated with environmental conditions, antagonistic trophic interactions can amplify habitat-filtering effects and thus lead to community clustering (Fine et al. 2006). Conversely, pathogens can increase competitive exclusion and thereby promote trait overdispersion (Gilbert and Webb 2007). Mutualistic interactions are also important in shaping communities (Bascompte and Jordano 2007). Pollinators shared among closely related plant species can, for example, increase phylogenetic clustering (Sargent and Ackerly 2008) and plants in early succession stages can facilitate co-occurrence of distantly related species which often lead to trait overdispersion (Valiente-Banuet and Verdu 2007). The bias in current inference methods, focusing on habitat filtering and competition is thus somewhat surprising.

Expanding into *geographical space*, the spatial distribution of habitats in relation to the dispersal propensity of organisms drives metapopulation and metacommunity dynamics, which alongside local ecological processes (Fig. 1) structures both competitive and trophic communities (Hanski 1999; Holyoak et al. 2005). Asynchrony in metapopulation dynamics and spatial dynamics in local extinction and recolonization of habitats can prevent extinctions (Hanski 1999; Holyoak et al. 2005). Such dynamics can be driven by multiple ecological mechanisms, such as different competitive advantages in different patches (Chesson 2000b; Chesson 2000a), competition-colonization trade-offs (Tilman 1994) and density-dependent predation (Holt 1993). Metacommunity dynamics (Fig. 1) can, however, also lead to extinctions, decrease the food-chain length and ultimately less diverse communities (Holt 1997). One can speculate that metapopulation dynamics that lead to species persisting where they would otherwise go extinct due to habitat filtering would render less community clustering, and that metapopulation dynamics that counteract competitive exclusion would increase community clustering. Such speculation is, however, difficult to confirm, as most current process-inference techniques do not consider metacommunity dynamics.

On *longer time scales* evolutionary processes can become important for the assembly of local communities (Fig. 1) (Urban and Skelly 2006). The absolute time on which this occurs is case dependent, and ecological and evolutionary time scales can overlap (Cortez and Ellner 2010). Hence, a mix of ecological and evolutionary processes assembles communities (Ellner et al. 2011). Knowledge of such eco-evolutionary processes and their effects on community structure is constantly increasing as, for example, theory explains how species adapt gradually according to selection gradients in a fitness landscape defined by the abiotic environment, resource availability and the traits and abundances of interacting species (see e.g., Brännström et al. 2013). Many empirical studies also demonstrate the importance of evolutionary processes at local spatial scales and character displacement due to competition, for example in Darwin’s finches (Schluter et al. 1985), may be quite common (Keller and Seehausen 2012). Furthermore, predation-induced trait divergence (Reznick et al. 2008; Zeller et al. 2012) can lead to decreased community clustering (Prinzing et al. 2008). Despite the evidence for eco-evolutionary processes being important, evolutionary processes are, however, poorly integrated into current inference techniques (Emerson and Gillespie 2008; Pausas and Verdu 2010).

*At larger spatial and longer temporal scales* (Fig. 1) the “evolving metacommunity” framework becomes relevant (Urban 2011; Mittelbach and Schemske 2015). This framework takes into account spatial variation in abiotic conditions and resource availability as well as dispersal and sequential colonization of species into a local community. A type of “race” between ecological (e.g. colonization) and evolutionary (e.g. local adaptation) processes occurs. Species can colonize a local community, adapt to novel conditions, and monopolize niche space before subsequent species invade (Urban and De Meester 2009). Conversely, invasion of well-adapted species can constrain evolutionary processes as niche space can be filled by well-adapted species, not through local adaptation(Urban et al. 2012). This “race” between ecological and evolutionary processes determines community and metacommunity structure and can be detected by, for example, phylogenetic structure analysis (Pontarp et al. 2012). Nevertheless, much-needed knowledge of the assembly processes and structure of spatially distributed evolving communities that also includes trophic interactions is lacking, though see Urban et al. (2008) for some conceptual examples.

## The case for inclusive and mechanistic process inference

Despite the complex nature of community assembly, inference methods often aim to infer about one or few processes, assuming the absence or at least no important influence of all others (Table 1; see Online Appendix 1 for a detailed review of the most common methods and their limitations). These inference techniques have been praised and criticized (see also Table 2 and review in Online Appendix 1) and calls for more inclusive and mechanistic approaches have been made (Mittelbach and Schemske 2015). Research aimed at addressing such calls exist, including the use of multiple existing inference techniques on the same data (Blois et al. 2014). Others aim at extensions and improvement of existing methods (Helmus et al. 2007; Leibold et al. 2010; De Bie et al. 2012). Such attempts, although necessary, remain associated with many of the challenges described in Table 2.

In line with advances in other fields, such as macroecology (D’Amen et al. 2015; Cabral et al. 2017) and ecological forecasting (Niu et al. 2014; Urban et al. 2016), we suggest that rather than using and developing existing inference techniques, a more general conceptual and flexible methodological inference framework should be adopted, coupling more realistic models with appropriate methods for fitting them to observed patterns. We present a generic eco-evolutionary and trait-based model as an example, coupled with Bayesian methods including model formulation, model fitting, and model improvement as a unified process inference approach (Fig 2). The framework can make use of *a priori* knowledge about the biological system studied, and although general, it is flexible enough to explain case-specific conditions. Different types of data can be utilized and the modeled mechanistic detail can be adjusted in accordance with different ecological realities, data types and data availability. Improvement of the inference is facilitated through quantitative evaluation of the inference quality and reliability.

**Fig. 2.**
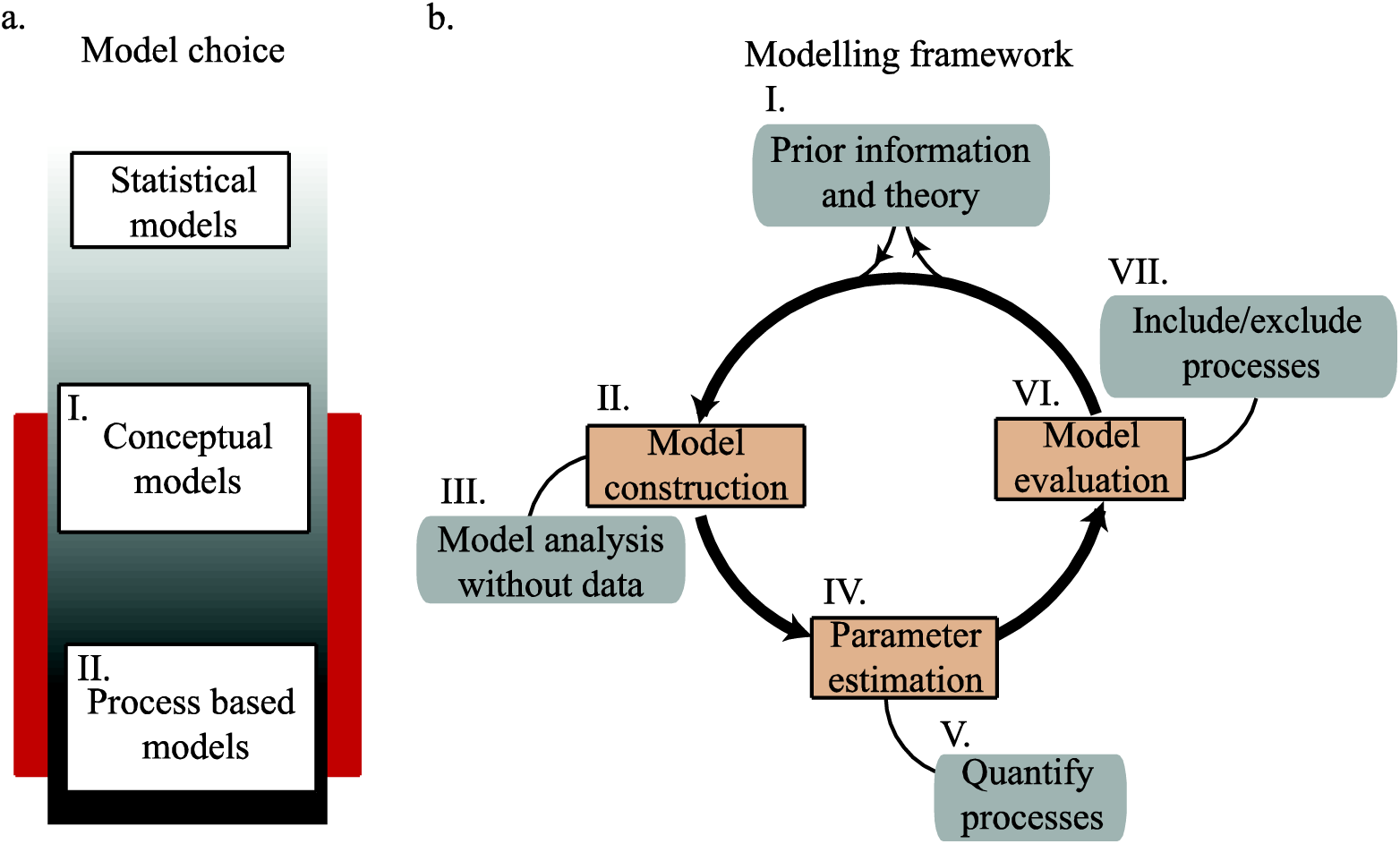
Illustration of the proposed process inference approach which involves three essential steps that feed into each other: 1) model construction, 2) model fitting and parameter estimation, and 3) model selection and model validation. For mechanistic process inference across scales, models need to include multiple processes and mechanistic detail but at the same time be simple enough to be computationally tractable. Dynamic community models are often based on simple population dynamical models but extended to include mechanisms through the inclusion of trait-based dynamics and population-or individual-based implementation (red section in a). Prior knowledge and theory inform model construction, including what processes to include, how to implement the model and at what level of organization and mechanistic detail the model should operate (b, I, II). Before the model is used for inference theoretical model investigation can identify different processes that may give rise to similar patterns and thus may be difficult to distinguish between (b, III). Parameter estimation provides quantitative information on the processes that are modeled given the data (b, IV, V) and the model selection procedure guide the model construction and inclusion or exclusion of particular processes (b, VI, VIII) and thus points out significant and critical processes that create patterns in data. The approach is independent of different data types, but the data will inform model construction.

## Implementing mechanistic and inclusive approaches

### Developing eco-evolutionary models for inference

Model construction is the first essential step in the inference framework proposed here, and it requires knowledge of the natural history of the study system, experiments, as well as known theory (Fig. 2). With Figure 1 in mind, it also becomes obvious that multiple processes should be included in the models as well as some mechanistic detail of those processes. Data (e.g. diversity, size distributions or phylogenetic patterns) also dictate model construction as model output and data needs to be comparable in subsequent parameter estimation and model selection steps (Fig. 2). It follows, that for a community model to be useful as a general inference tool, it needs to include multiple processes, it should be flexible, and it should output multiple types of data.

Dynamic models, which underlies much of our current understanding of communities, can be suitable for inference as it involves well-established functional forms and computational tractability (Brännström et al. 2012; Urban et al. 2016) (Fig. 2). The models are often made mechanistic through trait-based ecological interactions (Dieckmann and Doebeli 1999; Doebeli and Dieckmann 2003; Heinz et al. 2009) and evolutionary dynamics (e.g. Geritz et al. 1998; Dieckmann and Doebeli 1999; Doebeli and Dieckmann 2003).

Countless dynamic models have been analyzed and provided insights into ecology, evolution, and their interaction. As an example, Roughgarden (1972) used a trait-based modeled of a competitive community to study species co-existence. Evolutionary mechanisms such as mutation rates and mutation sizes have been studied in relation to trait evolution (Dieckmann and Law 1996). Others have implemented evolutionary mechanisms in models of co-evolution and trophic-community assembly (Ripa et al. 2009; Brännström et al. 2011). Spatial contingencies have been considered (Pontarp et al. 2015) and age- and stage structured populations, environmental and demographic stochasticity and variation in spatial structure can be included (Brännström et al. 2012).

Few studies have, however, utilized dynamic modeling for inference and none of the models presented above are suitable for inference in general as they are specifically designed to answer specific scientific questions. Dynamic eco-evolutionary models can, however, be constructed in a general and flexible way, fitting the requirements for inference. For the sake of argument, we implement such a model and we discuss its utility below.

We base our model on the generalized Lotka–Volterra (GLV) equations (Case 2000) extended into geographical space (Levin 1974). See Figure 3 for an illustration of the model and its initial values in our examples and see Appendix 2 for a detailed formulation of our model and description of the numerical implementation. Omitting space, for now, the per capita growth of *n* prey populations and *m* predator is formulated as:

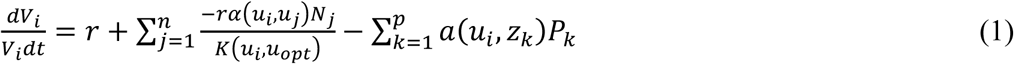

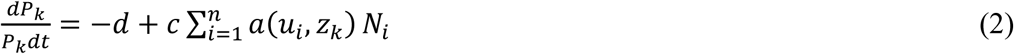

for *i* =1 to *n, k* =1 to *m* and where *V*_*i*_ and *P*_*k*_ denote prey and predator population size respectively. The parameter *r* and *d* is the intrinsic prey growth rate and the predator death rate, respectively. The functions on the right-hand side of equations 1 and 2 are trait dependent functions:

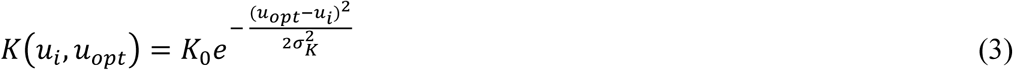

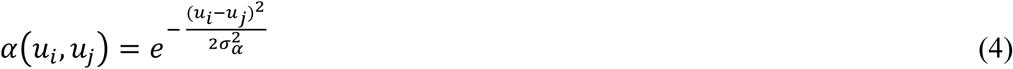

and

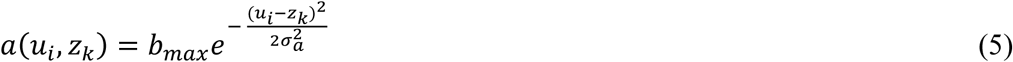

where *K*(*u*_*i*_, *u*_*opt*_) represents the carrying capacity for a monomorphic population *i* of prey individuals with trait value *u*_*i*_ in a habitat characterized by a resource distribution with its peak resource availability at the point *u*_*opt*_. It follows that the resource availability declines symmetrically as *u* deviates from *u*_*opt*_ according to the width of the resource distribution (*σK*). The interaction, *α*(*u*_*i*_,*u*_*j*_), between a prey population *i* (defined by its trait *u*_*i*_) and its competitor populations *j* (defined by their traits *u*_*j*_) is modeled in a similar way, through a Gaussian function. Here, we standardize the competition coefficients so that, for a focal population *i, α*_*ii*_ =1 and 0 < *α*_*ij*_<1 (*u*_*i*_≠*u*_*j*_). σ_α_ determines the degree of competition between individuals given certain utilization traits and can thus be viewed as the niche width of the prey. Equation 5 models the interaction, *a*(*u*_*i*_,*z*_*k*_), between a focal predator population *k* with trait value *z* and a prey population *i* with trait value *u*. The parameter *b*_*max*_ denotes the maximum attack rate obtained when *u*_*i*_=*z*_*k*_ and this rate then falls of symmetrically as *u*_*i*_ deviates from *z*_*k*_ according to a Gaussian function with variance *σ*_*a*_. Similar to the σ_α_ parameter, *σ*_*a*_ can be viewed as the niche width of the predator.

**Fig 3.**
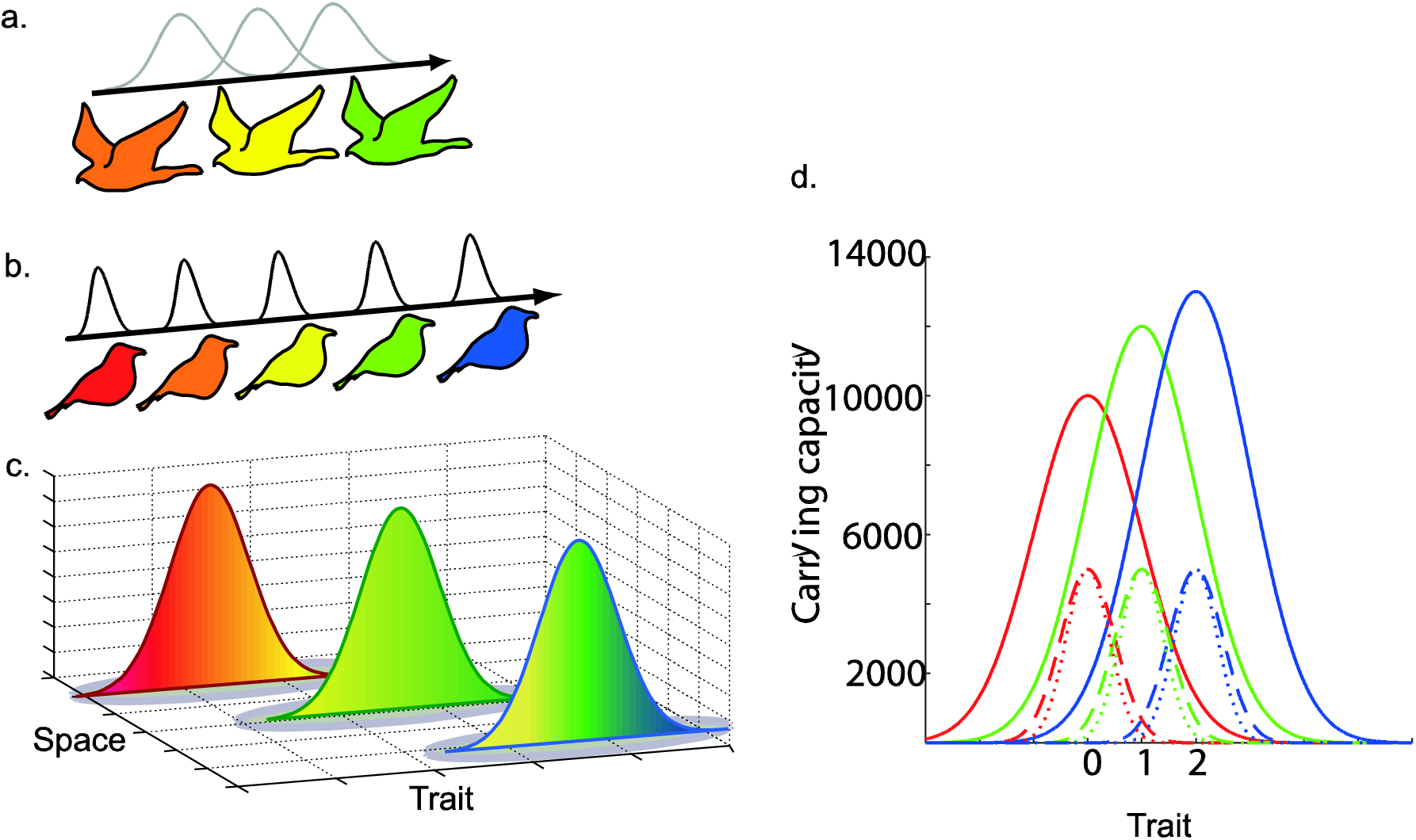
Illustration of the model (a-c) and initial conditions for the most complex model scenario presented this paper (d). Predators (a) and prey (b) can coexist and disperse between three habitats defined by their resource distributions (c). Species and resources are distributed in trait space (color gradient) and consumption is dictated by consumer-resource trait matching. As an example, red prey are optimized for utilizing red parts of the resource distribution and green predators are optimized to consume green prey. Note that in this illustration the blue prey has no predator. Competition between species is dictated by their niche width (black and gray Gaussian kernels). Large overlap between niche kernels indicates high competition and predation pressure, respectively. This model is flexible in the way the model can be initiated. In the most complex case, presented here, we initiate the model with three habitats each with their own resource distribution (solid red, green and blue lines in d). Maximum carrying capacity in each habitat is set to 10000, 12000, and 13000 and the peak of the distributions are situated at trait value 0, 1, and 2, respectively. We initiate the system with three prey and three predators (one in each habitat) with trait values equal to the resource distribution peaks. Prey and predators have the same trait value but we set niche widths for prey (dashed lines) to be slightly wider than predator niche width (dotted lines). Color coding denotes habitats and niche kernels are coded according to where the species occurs initially.Note that the y-axis has a direct association with the niche kernels in d.

We expand the non-spatial model described above into distinct patches or habitats distributed in space by implementing our model with a matrix formulation with vectors containing values for each population in each habitat, a community matrix that defines ecological interactions, and a dispersal matrix (Fig. 3 and Appendix 2). A fixed proportion of all local populations disperse between adjacent habitats. Furthermore, we follow an adaptive dynamics approach for the evolutionary dynamics (Geritz et al. 1998). In its full complexity, the model includes intra- and interspecific competition, trophic interactions, dispersal, trait evolution and in some cases evolutionary branching (Fig 3). The model provides us with information about population dynamics, equilibrium population sizes and trait distributions for each evolutionary step (Fig 4). Populations can also be assigned a species identity using, for example, a trait-based speciation definition (see also Pontarp et al. 2012; Pontarp et al. 2015). By registering the time and origin of all diversification events as well as trait distributions and abundance throughout evolutionary history we have all the information required to follow trait evolution, diversity, and phylogenetic and phenotypic community structure.

**Fig 4.**
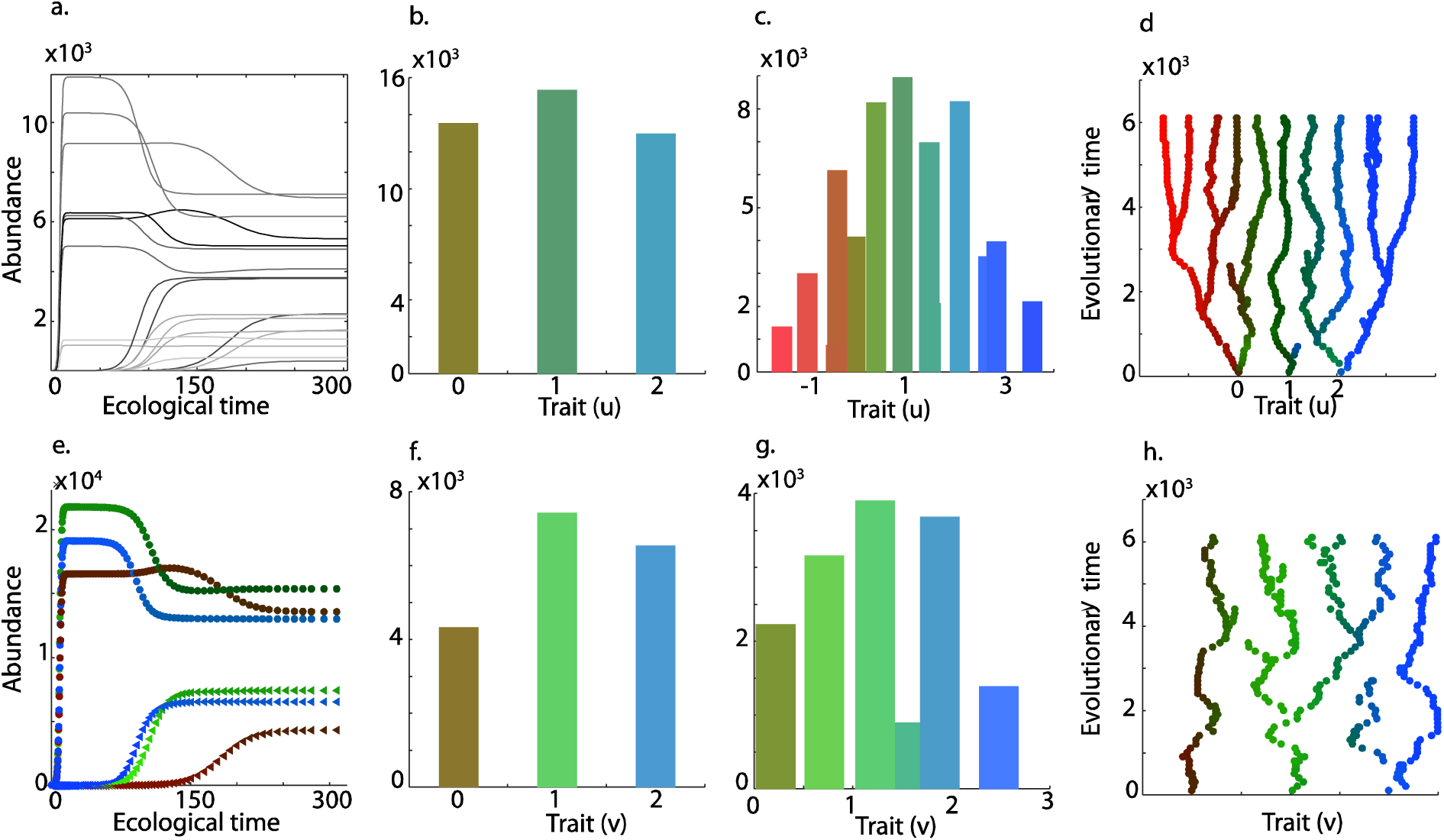
Representative data outputs from the model, including time series (a, e), trait distributions (n, c, f, g) and adaptive radiations (d, h) for prey (a, e, b, c, d) and predators (a, e, f, g, h). In simulations we first initiate the system (see initial conditions in Fig. 3) and we compute ecological dynamics and equilibrium. Here, we illustrate initiation with three prey and three predators distributed in three habitats (a). Panel e. contains the same data for prey (circles) and predator (triangles) populations as in panel a. but populations with similar morphs are summed across habitats. Color coding in e. illustrates spatial distribution. Pure red, green and blue denotes sole occupancy in habitat A, B, and C, respectively. Occupancy in multiple habitats are illustrated as a mix of colors proportional to the spatial distribution. Trait distributions at equilibrium for the initiated community are shown for the initiated community of three prey (b) and three predators (f) distributed in space. The spatial distribution and trait distributions of prey (c) and predators (g) evolved through adaptive radiations of co-evolving prey (d) and predators (h). Parameters for this simulation are: *K*_*0,*A_=10000; *K*_*0,*B_=12000; *K*_*0,*C_=13000; *u*opt,A = 0; *u*_opt,B_ = 1; *u*_opt,C_ = 2; *σK* =1; *σ*_*α*_ =0.5; *σ*_*a*_ =0.4; *r* = 1; *d* = −0.2; *cp* = 0.3;b_max_ = 0.0001; *M* = 0.05; *P*_mut.prey_ = 0.01; *P*_mut.pred_ = 0.1; *σ*_mut,prey_ =0.02; *σ*_mut,pred_ =0.03; with initial conditions *u*_*1*_ = 0; *u*_*2*_ = 2; *u*_*3*_ = 3; *ν*_*1*_ = 0; *ν*_*2*_ = 2; *ν*_*3*_ = 3.

### Tailoring mechanistic models to specific systems

Our model, as it is presented above and in Appendix 2, includes multiple processes and it can produce different types of data output (Figs 3-4). Our model can thus be used as an inference tool for complex systems, by searching for and finding distributions of parameter values (and therefore processes signs and strengths) that give the best correspondence between the model output and observations.

Unfortunately, increases in model complexity are accompanied by several challenges. Complex models tend to be difficult to interpret and the parameter estimation becomes increasingly computationally expensive and data demanding. Recent estimation techniques were developed with such challenges in mind (see below), but too complex models can become intractable and separating model structure error from parameter error becomes problematic (Keenan et al. 2011). Thus, the model should be made as simple as possible and still provide adequate information about the modeled community (May 2004). This can be accomplished by evaluating the data at hand and through prior knowledge of the ecology, evolution and natural history of the study system.

Prior knowledge of the study system may suggest that in some cases a relatively simple model is sufficient. A purely ecological model may, for example, be adequate for newly established communities of organisms with low evolutionary potential (e.g., low phenotypic/genotypic variation, low mutation rate, or low population sizes). In such cases, the model presented above can be reduced to an ecological community model, outputting time series data and trait distribution data only (Fig. 4 a). This is can be done by introducing species to a local community and allowing the community to assemble through ecological processes only, by omitting dispersal and the evolutionary algorithm altogether (Pontarp and Petchey 2016). For old communities or fast-evolving organisms (e.g., microbes) the full eco evolutionary model may be more appropriate, with or without the spatial component. Similarly, space may be omitted for largely sessile organisms in largely closed communities, while the inclusion of the spatial component of the model may be best for dispersing organisms and more open communities.

### Quantifying assembly processes through parameter estimation

Let us assume that prior information is available and data availability has guided us in our manipulation of the model such that we are relatively confident that we are modeling the correct processes. Now, model fitting and parameter estimation can provide information that is rarely provided by “traditional” inference approaches. Estimates of parameter distributions provide quantitative information about the processes with which the parameters are associated. By comparing estimates among parameters, or by sensitivity analysis, the relative strength and importance of different processes can be evaluated. Furthermore, the covariance between parameter distributions can inform about dependencies and redundancies between processes.

There are many methods for estimating model parameters and different approaches are appropriate for various types of models and data (Sokal and Rohlf 1995; Burnham and Anderson 2002; Hartig et al. 2011). These methods are reviewed elsewhere (reviewed in Raupach et al. 2005; Williams et al. 2009; Luo et al. 2011). Different fitting techniques will likely be preferred, depending on what type of processes are modeled and data availability. In simple cases where space and evolution are omitted, minimizing the sum of squared residuals may, for example, be preferred. Rather than review all possible fitting techniques for different model scenarios, we discuss issues that may arise when the model is complex and multiple data sources are available.

Approximate Bayesian Computation (ABC) is a model fitting technique with promise for overcoming the difficulty in fitting complex models to diverse data, and it could be the first option in such situations (Sisson et al. 2007; Beaumont 2010; Csillery et al. 2010; Blum et al. 2013). ABC takes priors for each model parameter as input, simulates data using the model and evaluates the distance (often Euclidian distance) between model output and data through a set of summary statistics (Fearnhead and Prangle 2012). The search of parameter space for the best performing parameters given data can be accomplished using global optimization techniques such as Kalman filters (Kalman 1960), Markov chain Monte Carlo (Gao et al. 2011) or Sequential Monte Carlo (Sisson et al. 2007). Posterior distributions for the parameters are approximated by rejecting or accepting parameter combinations through some distance threshold evaluated on the distance of summary statistics between model output and data. The approach consequently does not rely on computing the likelihood of the model given data as is done by more traditional frequentist or Bayesian fitting techniques. We thus view ABC as having great promise for parameter estimation and thereby process inference with complex process-based models.

### Inferring processes through model selection

Selecting among alternative model structures is the final essential step of inference. By iterating the model manipulation and model fitting steps, each time evaluating an increasingly complicated model, it is possible to circumvent potential problems of using an overly complex model from the start.

First, and before the models are fitted to real data, a theoretical model investigation can identify different processes that may give rise to similar patterns. If the models tell us that two processes give similar community patterns, it will be difficult to distinguish between those processes and additional information or even experimentation may be needed for successful inference. Second, it is possible to evaluate the intrinsic properties of the model versions and fitting techniques by testing them on simulated data produced by known parameter values. If the correct parameter values cannot be retrieved from simulated data, even though the model that underlie the patterns is known, correct inference on non-simulated (real) data, using that model and fitting technique, is unlikely. Third, while fitting models to data (Fig. 2) one can evaluate a model that includes fewer ecological processes against models that include more processes. The models are evaluated concerning how well they represent the data (Chivers et al. 2014), thus guiding the inclusion or exclusion of particular processes, and providing inference about processes.

Model selection is relatively straightforward when the models have the same number of parameters; goodness-of-fit can guide the selection. When the models have different numbers of parameters, other model selection techniques can be used. The most widely used are a suite of information criteria rooted in information theory (Akaike 1974). Other model selection criteria are, however, also possible. Again, ABC is a useful approach for complex models (Toni et al. 2009). The model selection procedure is based on the same general ABC principles presented above, except that the summary statistics is defined somewhat differently (Prangle et al. 2014). The output from the ABC model selection is focused on acceptance/rejection ratio between models rather than posterior parameter distributions (Toni et al. 2009; Liepe et al. 2014).

## Discussion

Ecological communities are complex, with diverse processes and actors (e.g. Urban and Skelly 2006; Vellend 2010; Urban et al. 2012; Mittelbach and Schemske 2015) and it is clear that several of the current inference techniques are too simplistic (Emerson and Gillespie 2008; Cavender-Bares et al. 2009; Vamosi et al. 2009; Cadotte et al. 2010; Pausas and Verdu 2010; Mouquet et al. 2012; Adler et al. 2013). A novel, more mechanistic, more inclusive, and more unified approach for future assembly-process inference techniques is desirable as this will allow us to infer the causal link between multiple processes and community patterns (Mittelbach and Schemske 2015), rather than focusing on less informative phenomenological/statistical relationships. We identify Bayesian analyses of model formulation, model fitting, and model improvement as a unified process-inference approach (Fig 2).

Approaches, similar to the ones presented above, have been suggested for predicting community response to environmental change (D’Amen et al. 2015; Urban et al. 2016). Furthermore, in ecological forecasting, a set of *ad hoc* models are often constructed and the best performing model is used for prediction (Luo et al. 2011; Niu et al. 2014; Urban et al. 2016). Although the approaches have not been synthesized for process-inference explicitly before, inference does, however, seem to be moving in the proposed direction. As an example, work on annual plants and parameterized models of competitor dynamics provides an understanding of how patterns of species coexistence are related to phylogenetic (Godoy et al. 2014) and phenotypic (Kraft et al. 2015) similarity. Massie et al. (2010) modeled trophic interactions and structured populations to infer drivers of community dynamics from phytoplankton population data. DeLong et al. (2014) inferred predator-prey interactions from microcosm experimental data to better understand ecological drivers of predator body size. On larger spatial scales Carrara et al. (2012) used a spatially explicit population model to infer processes from microcosm metacommunities. Furthermore, Yoshida et al. (2007) parameterized an evolutionary predator-prey model and infer interaction strength from community dynamics.

Studies that more explicitly use the proposed inference approach presented above also exist. Recently, van der Plas et al. (2015) published a modeling approach for estimating the relative importance of different community assembly processes. They used a trait-based (but not dynamic as described above) model and they simulated the assembly through processes like dispersal, habitat filtering and limiting similarity. They then fitted their model to community data using ABC, and by estimating parameters that are directly linked to the strength of the different processes, they inferred the relative strength of those processes. Jabot and Bascompte (2012) used a similar approach to contrast dispersal limitations and stochastic metacommunity dynamics against trophic interactions. They too used a simulation approach to assemble, in this case, network communities and they used ABC to parameterize the model given data. Ultimately they inferred how trophic interactions shape biodiversity.

May et al. (2013) went even further and used ABC for parameter estimation and model selection as an inference tool. They also used simulations, and they contrasted a metacommunity model, a mainland-island model and an island community model against each other. The best performing model, given vegetation survey data, was used to inferring the role of connectivity through seed dispersal among habitat patches for regional community dynamics. Although they do not present their work as an iterative framework including model construction, parameter estimation and model evaluation, the studies presented here are excellent examples where more or less mechanistic models were used for process inference.

The proposed framework is general and from the literature reviewed above, we conclude that several modeling approaches and statistical techniques can be used. The synthesis of the relatively simplistic and flexible nature of dynamical modeling in combination with the powerful and flexible ABC does, however, seem particularly suitable. Dynamic modeling is simple in the sense that it is based on simple, often phenomenological population dynamical models (Brännström et al. 2012). The “skeleton” of such simple models are then extended to include detailed mechanisms through the inclusion of trait-based dynamics, complex functional forms and through a population- or individual-based implementation. The trade-off between realism, computational costs, and model tractability can be monitored and controlled as the models gain in complexity. By iterating over model construction and model evaluation several times, each time evaluating an increasingly complex version of the model, the optimal model for the study system and data can be found. Any model that include several processes will, however, tend to be complex, computationally costly and likelihood functions are often intractable, leading to the need of powerful fitting and model selection techniques like ABC.

It is also important to emphasize the empirical side of inference, namely data to which the models are fitted. As noted above, data inform the models and is thus imperative for the model fitting (Urban et al. 2016; Cabral et al. 2017). Data also dictate model construction as the model output needs to be comparable with data (e.g. diversity, size distributions or phylogenetic patterns). Furthermore, data provide knowledge of the natural history of the study system that also informs model construction. *A priori* information of a particular system can narrow down the priors for ABC and thus facilitate parameter estimation by reducing the parameter space that needs to be searched in the optimization procedure. For certain systems, reasonable parameter values may already be available in the literature. In other cases, it might be possible to measure some parameters in independent studies. As an example, DeLong et al. (2014) conducted separate experiments to estimate functional responses before they fitted a full predator-prey model to protist microcosm data. Similarly, Kraft et al. (2015) constructed the functional form for their model following information provided by experiments before they inferred vital rates and pairwise competitive interactions. Kraft’s work again illustrates the importance of prior and separate sources of knowledge and high-quality data, and it highlights the importance of combining experimental work with inference from observations. The experimental work reviewed in this paper are examples of how mechanistic modeling and parameter estimation techniques combine to provide a better understanding of community assembly and dynamics in general as well as to enable better inference of community assembly processes from observed macroscopic patterns.

## Conclusion

We envision that process inference will continue to move away from simple statistical and non-mechanistic inference techniques for approaches with a constant flow of information between experimental and field data, model construction, parameter estimation, and model selection. This way the challenges associated with inference (Table 2), may be avoided and the full complexity of community and structure can be more and more considered and understood. The transformation in inference approach will come with technical challenges as well as increased demands on data available from natural systems, computational power, and experimental progress. Many of these difficulties are, however, already identified and to some extent resolved in other fields and thus ready to be put into action in a more formal way also for inferring processes of community assembly from macroscopic patterns.

## Acknowledgements

We thank Jonathan Levine, Florian Altermatt and Miguel Verdú for comments and suggestions that improved the initial version of this paper.

## References

Ackerly, D. D., D. W. Schwilk, and C. O. Webb. 2006. Niche evolution and adaptive radiation: Testing the order of trait divergence. Ecology 87:50–61.

Adler, P. B., A. Fajardo, A. R. Kleinhesselink, and N. J. B. Kraft. 2013. Trait-based tests of coexistence mechanisms. Ecology Letters 16:1294–1306.

Akaike, H. 1974. A new look at the statistical model identification. IEEE Transactions on Automatic Control 19:716–723.

Alto, B. W., J. Malicoate, S. M. Elliott, and J. Taylor. 2012. Demographic consequences of predators on prey: Trait and density mediated effects on mosquito larvae in containers. Plos One 7:1–8.

Bascompte, J., and P. Jordano. 2007. Plant-animal mutualistic networks: The architecture of biodiversity. Annual Review of Ecology Evolution and Systematics 38:567–593.

Beaumont, M. A. 2010. Approximate Bayesian Computation in Evolution and Ecology. Annual Review of Ecology, Evolution, and Systematics, Vol 41 41:379–406.

Blois, J. L., N. J. Gotelli, A. K. Behrensmeyer, J. T. Faith, S. K. Lyons, J. W. Williams, K. L. Amatangelo et al. 2014. A framework for evaluating the influence of climate, dispersal limitation, and biotic interactions using fossil pollen associations across the late Quaternary. Ecography 37:1095–1108.

Blomberg, S. P., T. Garland, and A. R. Ives. 2003. Testing for phylogenetic signal in comparative data: Behavioral traits are more labile. Evolution 57:717–745.

Blum, M. G. B., M. A. Nunes, D. Prangle, and S. A. Sisson. 2013. A comparative review of dimension reduction methods in Approximate Bayesian Computation. Statistical Science 28:189–208.

Borcard, D., P. Legendre, and P. Drapeau. 1992. Partialling out the spatial component of ecological variation. Ecology 73:1045–1055.

Brännström, A., J. Johansson, N. Loeuille, N. Kristensen, T. A. Troost, R. H. R. Lambers, and U. Dieckmann. 2012. Modelling the ecology and evolution of communities: a review of past achievements, current efforts, and future promises. Evolutionary Ecology Research 14:601–625.

Brännström, å., J. Johansson, and N. von Festenberg. 2013. The hitchhiker’s guide to adaptive dynamics. Games 4:304–328.

Brännström, A., N. Loeuille, M. Loreau, and U. Dieckmann. 2011. Emergence and maintenance of biodiversity in an evolutionary food-web model. Theoretical Ecology 4:467–478.

Burnham, K. P., and D. R. Anderson. 2002, Model selection and multimodel inference: A practical Information-theoretic approach. New York, Springer.

Cabral, J. S., L. Valente, and F. Hartig. 2017. Mechanistic simulation models in macroecology and biogeography: state-of-art and prospects. Ecography 40.

Cadotte, M. W., T. J. Davies, J. Regetz, S. W. Kembel, E. Cleland, and T. H. Oakley. 2010. Phylogenetic diversity metrics for ecological communities: integrating species richness, abundance and evolutionary history. Ecology Letters 13:96–105.

Carrara, F., F. Altermatt, I. Rodriguez-Iturbe, and A. Rinaldo. 2012. Dendritic connectivity controls biodiversity patterns in experimental metacommunities. Proceedings of the National Academy of Sciences of the United States of America 109:5761–5766.

Case, T. J. 2000, An illustrated guide to theoretical ecology. Oxford New York, Oxford University Press, Inc.

Cavender-Bares, J., K. H. Kozak, P. V. A. Fine, and S. W. Kembel. 2009. The merging of community ecology and phylogenetic biology. Ecology Letters 12:693–715.

Chesson, P. 2000a. General theory of competitive coexistence in spatially-varying environments. Theoretical Population Biology 58:211–237.

Chesson, P.. 2000b. Mechanisms of maintenance of species diversity. Annual Review of Ecology and Systematics 31:343–366.

Chivers, C., B. Leung, and N. D. Yan. 2014. Validation and calibration of probabilistic predictions in ecology. Methods in Ecology and Evolution 5:1023–1032.

Cortez, M. H., and S. P. Ellner. 2010. Understanding rapid evolution in predator-prey interactions using the theory of fast-slow dynamical systems. American Naturalist 176:E109–E127.

Csillery, K., M. G. B. Blum, O. E. Gaggiotti, and O. Francois. 2010. Approximate Bayesian Computation (ABC) in practice. Trends in Ecology & Evolution 25:410–418.

D’Amen, M., C. Rahbek, N. E. Zimmermann, and A. Guisan. 2015. Spatial predictions at the community level: from current approaches to future frameworks. Biological Reviews:/a-n/a.

De Bie, T., L. De Meester, L. Brendonck, K. Martens, B. Goddeeris, D. Ercken, H. Hampel et al. 2012. Body size and dispersal mode as key traits determining metacommunity structure of aquatic organisms. Ecology Letters 15:740–747.

DeLong, J. P., T. C. Hanley, and D. A. Vasseur. 2014. Predator-prey dynamics and the plasticity of predator body size. Functional Ecology 28:487–493.

Dieckmann, U., and M. Doebeli. 1999. On the origin of species by sympatric speciation. Nature 400:354–357.

Dieckmann, U., and R. Law. 1996. The dynamical theory of coevolution: A derivation from stochastic ecological processes. Journal of Mathematical Biology 34:579–612.

Doebeli, M., and U. Dieckmann. 2003. Speciation along environmental gradients. Nature 421:259–264.

Dray, S., and P. Legendre. 2008. Testing the species traits-environment relationships: The fourth-corner problem revisited. Ecology 89:3400–3412.

Ellner, S. P., M. A. Geber, and N. G. Hairston. 2011. Does rapid evolution matter? Measuring the rate of contemporary evolution and its impacts on ecological dynamics. Ecology Letters 14:603–614.

Emerson, B. C., and R. G. Gillespie. 2008. Phylogenetic analysis of community assembly and structure over space and time. Trends in Ecology & Evolution 23:619–630.

Fearnhead, P., and D. Prangle. 2012. Constructing summary statistics for approximate Bayesian computation: semi-automatic approximate Bayesian computation. Journal of the Royal Statistical Society Series B-Statistical Methodology 74:419–474.

Fine, P. V. A., Z. J. Miller, I. Mesones, S. Irazuzta, H. M. Appel, M. H. H. Stevens, I. Saaksjarvi et al. 2006. The growth-defense trade-off and habitat specialization by plants in Amazonian forests. Ecology 87:S150–S162.

Gao, C., H. Wang, E. S. Weng, S. Lakshmivarahan, Y. F. Zhang, and Y. Q. Luo. 2011. Assimilation of multiple data sets with the ensemble Kalman filter to improve forecasts of forest carbon dynamics. Ecological Applications 21:1461–1473.

Geritz, S. A. H., E. Kisdi, G. Meszena, and J. A. J. Metz. 1998. Evolutionarily singular strategies and the adaptive growth and branching of the evolutionary tree. Evolutionary Ecology 12:35–57.

Gilbert, G. S., and C. O. Webb. 2007. Phylogenetic signal in plant pathogen-host range. Proceedings of the National Academy of Sciences of the United States of America 104:4979–4983.

Godoy, O., N. J. B. Kraft, and J. M. Levine. 2014. Phylogenetic relatedness and the determinants of competitive outcomes. Ecology Letters 17:836–844.

Gotelli, N. J., M. J. Anderson, H. T. Arita, A. Chao, R. K. Colwell, S. R. Connolly, D. J. Currie et al. 2009. Patterns and causes of species richness: a general simulation model for macroecology. Ecology Letters 12:873–886.

Hanski, I. 1999, Metapopulation Ecology. Oxford, Oxford University Press.

Hartig, F., J. M. Calabrese, B. Reineking, T. Wiegand, and A. Huth. 2011. Statistical inference for stochastic simulation models - theory and application. Ecology Letters 14:816–827.

Heinz, S. K., R. Mazzucco, and U. Dieckmann. 2009. Speciation and the evolution of dispersal along environmental gradients. Evolutionary Ecology 23:53–70.

Helmus, M. R., T. J. Bland, C. K. Williams, and A. R. Ives. 2007. Phylogenetic measures of biodiversity. American Naturalist 169:E68–E83.

Holt, R. D. 1993, Ecology at the mesoscale: The influence of regional processes on local communities. Chicago, Chicago University Press.—. 1997. Community modules. Multitrophic Interactions in Terrestrial Systems:333-350.

Holyoak, M., M. A. Leibold, and R. D. Holt. 2005, Metacommunities: spatial dynamics and ecological communities, University of Chicago Press.

Jabot, F., and J. Bascompte. 2012. Bitrophic interactions shape biodiversity in space. Proceedings of the National Academy of Sciences of the United States of America 109:4521–4526.

Kalman, R. E. 1960. A new approach to linear filtering and prediction problems. Journal of basic engineering 82:35–45.

Keenan, T. F., M. S. Carbone, M. Reichstein, and A. D. Richardson. 2011. The model-data fusion pitfall: assuming certainty in an uncertain world. Oecologia 167:587–597.

Keller, I., and O. Seehausen. 2012. Thermal adaptation and ecological speciation. Molecular Ecology 21:782–799.

Kot, M. 2001, Mathematical Ecology. Cambridge, Cambridge University Press.

Kraft, N. J. B., O. Godoy, and J. M. Levine. 2015. Plant functional traits and the multidimensional nature of species coexistence. Proceedings of the National Academy of Sciences of the United States of America:797–802.

Legendre, P., R. Galzin, and M. L. HarmelinVivien. 1997. Relating behavior to habitat: Solutions to the fourth-corner problem. Ecology 78:547–562.

Leibold, M. A., E. P. Economo, and P. Peres-Neto. 2010. Metacommunity phylogenetics: separating the roles of environmental filters and historical biogeography. Ecology Letters 13:1290–1299.

Levin, S. A. 1974. Dispersion and Population Interactions. American Naturalist 108:207–228.

Liepe, J., P. Kirk, S. Filippi, T. Toni, C. P. Barnes, and M. P. H. Stumpf. 2014. A framework for parameter estimation and model selection from experimental data in systems biology using approximate Bayesian computation. Nature Protocols 9:439–456.

Luo, Y. Q., K. Ogle, C. Tucker, S. F. Fei, C. Gao, S. LaDeau, J. S. Clark et al. 2011. Ecological forecasting and data assimilation in a data-rich era. Ecological Applications 21:1429– 1442.

MacArthur, R. H., and R. Levins. 1967. Limiting similarity convergence and divergence of coexisting species. American Naturalist 101:377–385.

Massie, T. M., B. Blasius, G. Weithoff, U. Gaedke, and G. F. Fussmann. 2010. Cycles, phase synchronization, and entrainment in single-species phytoplankton populations. Proceedings of the National Academy of Sciences of the United States of America 107:4236–4241.

May, F., I. Giladi, M. Ristow, Y. Ziv, and F. Jeltsch. 2013. Metacommunity, mainland-island system or island communities? Assessing the regional dynamics of plant communities in a fragmented landscape. Ecography 36:842–853.

May, R. M. 2004. Uses and abuses of mathematics in biology. Science 303:790–793.

Mayfield, M. M., and J. M. Levine. 2010. Opposing effects of competitive exclusion on the phylogenetic structure of communities. Ecology Letters 13:1085–1093.

Mittelbach, G. G., and D. W. Schemske. 2015. Ecological and evolutionary perspectives on community assembly. Trends in Ecology & Evolution 30:241–247.

Mouquet, N., V. Devictor, C. N. Meynard, F. Munoz, L. F. Bersier, J. Chave, P. Couteron et al. 2012. Ecophylogenetics: advances and perspectives. Biological Reviews 87:769–785.

Niu, S. L., Y. Q. Luo, M. C. Dietze, T. F. Keenan, Z. Shi, J. W. Li, and F. S. Chapin. 2014. The role of data assimilation in predictive ecology. Ecosphere 5.

Pausas, J. G., and M. Verdu. 2010. The jungle of methods for evaluating phenotypic and phylogenetic structure of communities. Bioscience 60:614–625.

Petchey, O. L. 2007. Effects of environmental variability on ecological communities: Testing the insurance hypothesis of biodiversity in aquatic microcosms. Impact of Environmental Variability on Ecological Systems 2:179–196.

Petchey, O. L., and K. J. Gaston. 2006. Functional diversity: back to basics and looking forward. Ecology Letters 9:741–758.

Pontarp, M., and O. L. Petchey. 2016. Community trait overdispersion due to trophic interactions: concerns for assembly process inference. Proceedings of the Royal Society B-Biological Sciences 283.

Pontarp, M., J. Ripa, and P. Lundberg. 2012. On the origin of phylogenetic structure in competitive metacommunities. Evolutionary Ecology Research 14:269–284.

Pontarp, M., J. Ripa. 2015. The biogeography of adaptive radiations and the geographic overlap of sister species. American Naturalist 186:565–581.

Prangle, D., P. Fearnhead, M. P. Cox, P. J. Biggs, and N. P. French. 2014. Semi-automatic selection of summary statistics for ABC model choice. Statistical Applications in Genetics and Molecular Biology 13:67–82.

Prinzing, A., R. Reiffers, W. G. Braakhekke, S. M. Hennekens, O. Tackenberg, W. A. Ozinga, J. H. J. Schaminee et al. 2008. Less lineages - more trait variation: phylogenetically clustered plant communities are functionally more diverse. Ecology Letters 11:809–819.

Raupach, M. R., P. J. Rayner, D. J. Barrett, R. S. DeFries, M. Heimann, D. S. Ojima, S. Quegan et al. 2005. Model-data synthesis in terrestrial carbon observation: methods, data requirements and data uncertainty specifications. Global Change Biology 11:378–397.

Reznick, D. N., C. K. Ghalambor, and K. Crooks. 2008. Experimental studies of evolution in guppies: a model for understanding the evolutionary consequences of predator removal in natural communities. Molecular Ecology 17:97–107.

Ricklefs, R. E. 2007. History and diversity: Explorations at the intersection of ecology and evolution. American Naturalist 170:56–70.

Ripa, J., L. Storlind, P. Lundberg, and J. S. Brown. 2009. Niche co-evolution in consumer-resource communities. Evolutionary Ecology Research 11:305–323.

Roughgarden, J. 1972. Evolution of niche width. American Naturalist 106:683–718.

Sargent, R. D., and D. D. Ackerly. 2008. Plant-pollinator interactions and the assembly of plant communities. Trends in Ecology & Evolution 23:123–130.

Schluter, D., T. D. Price, and P. R. Grant. 1985. Ecological Character Displacement in Darwin Finches. Science 227:1056–1059.

Sisson, S. A., Y. Fan, and M. M. Tanaka. 2007. Sequential Monte Carlo without likelihoods. Proceedings of the National Academy of Sciences of the United States of America 104:1760–1765.

Sokal, R. R., and F. J. Rohlf. 1995, The principles and practice of statistics in biological research. Stony Brook University, W.H. Freeman & Co.

Tilman, D. 1994. Competition and biodiversity in spatially structured habitats. Ecology 75:2– 16.

Toni, T., D. Welch, N. Strelkowa, A. Ipsen, and M. P. H. Stumpf. 2009. Approximate Bayesian computation scheme for parameter inference and model selection in dynamical systems. Journal of the Royal Society Interface 6:187–202.

Trisos, C. H., O. L. Petchey, and J. A. Tobias. 2014. Unraveling the interplay of community assembly processes acting on multiple niche axes across spatial scales. American Naturalist 184:593–608.

Urban, M. C. 2011. The evolution of species interactions across natural landscapes. Ecology Letters 14:723–732.

Urban, M. C., G. Bocedi, A. P. Hendry, J. B. Mihoub, G. Pe’er, A. Singer, J. R. Bridle et al. 2016. Improving the forecast for biodiversity under climate change. Science 353.

Urban, M. C., and L. De Meester. 2009. Community monopolization: local adaptation enhances priority effects in an evolving metacommunity. Proceedings of the Royal Society B-Biological Sciences 276:4129–4138.

Urban, M. C., L. De Meester, M. Vellend, R. Stoks, and J. Vanoverbeke. 2012. A crucial step toward realism: responses to climate change from an evolving metacommunity perspective. Evolutionary Applications 5:154–167.

Urban, M. C., M. A. Leibold, P. Amarasekare, L. De Meester, R. Gomulkiewicz, M. E. Hochberg, C. A. Klausmeier et al. 2008. The evolutionary ecology of metacommunities. Trends in Ecology & Evolution 23:311–317.

Urban, M. C., and D. K. Skelly. 2006. Evolving metacommunities: Toward an evolutionary perspective on metacommunities. Ecology 87:1616–1626.

Valiente-Banuet, A., and M. Verdu. 2007. Facilitation can increase the phylogenetic diversity of plant communities. Ecology Letters 10:1029–1036.

Vamosi, S. M., S. B. Heard, J. C. Vamosi, and C. O. Webb. 2009. Emerging patterns in the comparative analysis of phylogenetic community structure. Molecular Ecology 18:572– 592.

van der Plas, F., T. Janzen, A. Ordonez, W. Fokkema, J. Reinders, R. S. Etienne, and H. Olff. 2015. A new modeling approach estimates the relative importance of different community assembly processes. Ecology 96:1502–1515.

Vellend, M. 2010. Conceptual synthesis in community ecology. Quarterly Review of Biology 85:183–206.

Webb, C. O., D. D. Ackerly, M. A. McPeek, and M. J. Donoghue. 2002. Phylogenies and community ecology. Annual Review of Ecology and Systematics 33:475–505.

Wiens, J. J., D. D. Ackerly, A. P. Allen, B. L. Anacker, L. B. Buckley, H. V. Cornell, E. I. Damschen et al. 2010. Niche conservatism as an emerging principle in ecology and conservation biology. Ecology Letters 13:1310–1324.

Williams, M., A. D. Richardson, M. Reichstein, P. C. Stoy, P. Peylin, H. Verbeeck, N. Carvalhais et al. 2009. Improving land surface models with FLUXNET data. Biogeosciences 6:1341–1359.

Wilson, J. B., E. Weiher, and P. Keddy. 1999, Assembly rules in plant communities. Cambridge, Cambridge University Press.

Yoshida, T., S. P. Ellner, L. E. Jones, B. J. M. Bohannan, R. E. Lenski, and N. G. Hairston. 2007. Cryptic population dynamics: Rapid evolution masks trophic interactions. Plos Biology 5:1868–1879.

Zeller, M., K. Lucek, M. P. Haesler, O. Seehausen, and A. Sivasundar. 2012. Signals of predation-induced directional and disruptive selection in the threespine stickleback. Evolutionary Ecology Research 14:193–205.

